# Rapid Detection of Norovirus Using Paper-based Microfluidic Device

**DOI:** 10.1101/162396

**Authors:** Xuan Weng, Suresh Neethirajan

## Abstract

Noroviruses (NoV) are the leading cause of outbreak of acute gastroenteritis worldwide. A substantial effort has been made in the development of analytical devices for rapid and sensitive food safety monitoring via the detection of foodborne bacteria, viruses and parasites. Conventional analytical approaches for noroviruses suffer from some critical weaknesses: labor-intensive, time-consuming, and relatively low sensitivity. In this study, we developed a rapid and highly sensitive biosensor towards point-of-care device for noroviruses based on 6-carboxyfluorescein (6-FAM) labeled aptamer and nanomaterials, multi-walled carbon nanotubes (MWCNTs) and graphene oxide (GO). In an assay, the fluorescence of 6-FAM labeled aptamer was quenched by MWCNTs or GO via fluorescence resonance energy transfer (FRET). In the presence of norovirus, the fluorescence would be recovered due to the release of the 6-FAM labeled aptamer from MWCNTs or GO. An easy-to-make paper-based microfluidic platform made by nitrocellulose membrane was used to conduct the assay. The quantitative detection of norovirus virus-like particles (NoV VLPs) was successfully performed. A linear range of 0-12.9 μg/mL with a detection limit of 40 pM and 30 pM was achieved for the MWCNTs and GO based paper sensors, respectively. The results suggested the developed paper-based microfluidic device is simple, cost-effective and holds the potential of rapid in situ visual determination for noroviruses with remarkable sensitivity and specificity, which provides a new way for early identification of NoV and thereby an early intervention for preventing the spread of an outbreak.

## Introduction

Noroviruses (NoV) are the leading causes of acute viral gastroenteritis [1], as well as one of the leading causes of foodborne disease outbreaks and the most common cause of infectious gastroenteritis worldwide [2], clinical symptoms including abdominal pain, diarrhea, muscle ache, and mild fever [3]. NoV infection may be direct from person to person or transmitted through food and drinks (ready-to-eat foods, table water) and the outbreak of which is frequently occurred in community facilities such as hospitals and school cafeterias. Currently, NoV are usually detected after an outbreak on the suspected food with a range of methods, including enzyme-linked immu-nosorbent assay (ELISA) [4], western blot [5], and nucleic acid-based methods, such as reverse transcription polymerase chain reaction (RT-PCR) [6] etc. These methods usually require expensive reagents, specialized equipment, and skillful researchers to perform the analyses and long waiting time for the results. In addition, antibodies are usually used in the biomolecular recognition, however, antibodies are usually expensive, have limited shelf-life and false-positive issues may occur due to the cross-reactivity between the antibodies and non-target molecules. Since the NoV have high resistance to routine disinfection and low infection dose, early on-site detection is extremely important for the prevention of the transmission and effective controlling of the infection as well as the spread of an outbreak. In addition, NoV, in cases, can be pre-emptively identified to efficiently block and reduce the infection [7]. FDA of United States approved the first early detection NoV test in 2011. All of the aforementioned reasons urge the need for more rapid, accurate, and ultra-sensitive assays to detect potential NoV contamination in complex food matrices. Development of such a method for NoV associated with foodborne illness can be a powerful tool for the application in response to outbreak management.

To date, researchers have put numerous efforts in the development of rapid and efficient methods, and biosensor is one of the promising ways to provide alternative to conventional assays due to its features of rapid, simple, reproducible, and cost-effective. Hong et al. [8] reported an electrochemical biosensor for pre-emptive on-site NoV detection by using nanostructured gold electrode conjugated with a NoV capturing agent concanavalin A. The biosensor was able to detect NoV in a concentration range of 10^2^ and 10^6^ copies/mL within 1 hour. Hwang et al. [9] reported a label-free electrochemical biosensor for human NoV detection using affinity peptide and a limit of detection (LoD) of 7.8 copies/mL was achieved. Although electrochemical biosensors are able to provide high sensitivity detection, the relative high price make them not suitable to be used in disposable chips for on-site detection [10]. Lateral-flow assays (LFAs) have been well developed and widely used in point-of-care testing (POCT), which have also been applied for virus detection [11, 12]. The main drawbacks of most of LFAs are the low sensitivity and the yes/no results. The introduction of nanomaterial contributes to the enhancement of sensitivity. Hagström et al. [13] developed a LFA for the detection of NoV by using nanoparticle reporters and a LoD of 10^7^ virus-like particles per microliter was obtained.

Paper-based microfluidic devices, as alternative to the traditional microfluidic devices made by PDMS, glass, silicon or other polymers, find their applications in biochemical, health diagnosis and food safety fields due to its unique advantages, including the low cost, easy-to-fabricate process and the simplicity [14, 15]. In addition, paper-based microfluidic devices are powerless due to the capillary action of the hydrophobic channel. Currently, wax printing technique is being used to fabricate the paper-based microfluidic devices.

Therefore, we developed a simple, low-cost and easy-to-make paper-based microfluidic device with multi-detection capability for determination of virus contaminants in food. NoV was used as the testing model. Carbon nanostructures usually have a wide range of absorption spectrum overlapping with the fluorescence spectra of various fluorophores [16], which allows FRET between them. The paper-based microfluidic device utilized FRET between nanomaterials and the virus probe functionalized fluorophore to quantitatively determine the concentration of the sample. In the present work, 6-carboxyfluorescein (6-FAM) labeled norovirus aptamer associated with MWCNTs or GO was employed as the “probe” to sensitively sense the presence of NoV with high specificity. MWCNTs consist of multiple rolled layers (concentric tubes) of graphitic sheets [17] while GO is a single-atom-thick two-dimensional carbon nanomaterial and both of them are efficient quenchers to various fluorophores [18, 19]. In our study, the efficiency of two kinds of nanomaterials namely, MWCNTs and GO, as the fluorescence quencher were investigated. In addition, the sensing probe was operated on an easy-to-make paper-based microfluidic device which is inexpensive and can be easily fabricated by a paper puncher within a minute. Such paper-based microfluidic device is power free because sample can be driven by the capillary force, and can be easily made within 1 min. The design of the paper-based microfluidic device allows for multiplex detection (up to 6). Another advantage of using paper substrate is the avoidance of complicated processes of probe immobilization and surface modification which can be achieved through the physical absorption. Schematic illustration in the Fig. 1a and 1b shows the main principle of this sensor, as an example, only the MWCNTs is illustrated. The principle of the sensing is based on the FRET between the fluorophores and nanostructure quenchers. The conformational change of the aptamer associated with the distance changes between the fluorophores and quenchers leads to a measurable fluorescence intensity. A ready-to-use device was made by loading the complex “probe” consisting of nanomaterial and target specific aptamer with fluorescence labelling onto the reaction wells and dries out. In an assay, sample was loaded onto the sample well followed by diffusion to the reaction wells immediately and bind with the aptamers, leading to the recovery of the fluorescence. The recovered fluorescence intensity was then used to quantitatively determine the target concentration in the sample.

**Fig. 1.**
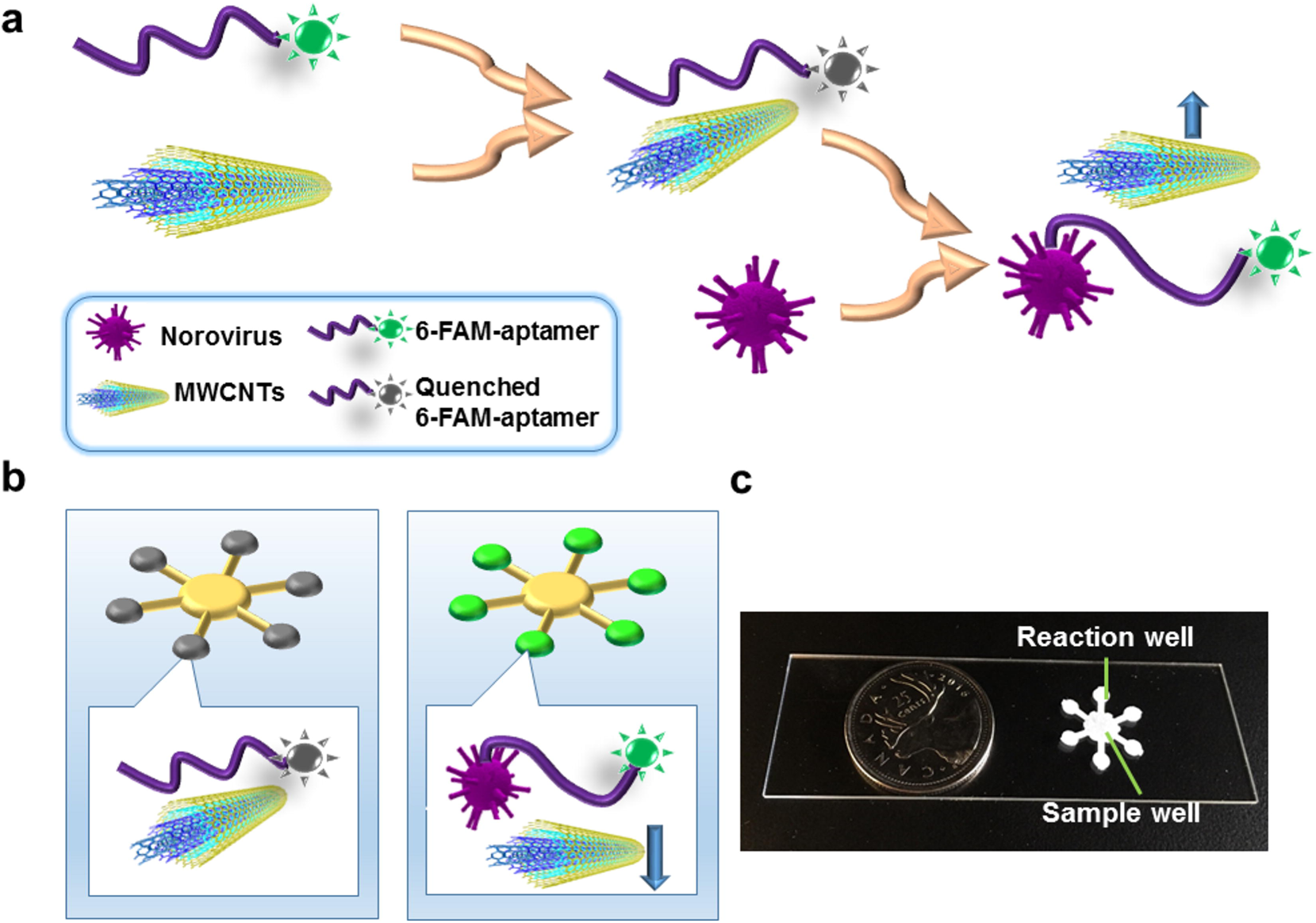
**a** Schematic illustration of turn-on sensor based on MWCNTs and 6-FAM functionalized aptamer, illustrating the principle of the NoV detection based on the nanomaterial MWCNTs (GO), Nov aptamer and NoV VLPs. The fluorescence of the NoV aptamer would be quenched when they were mixed and coated on the paper surface due to the FRET process between the probe and GO. In the presence of the NoV, the aptamer would bond to the protein because of the stronger association constant between them, resulting in the fluorescence revocery. **b** Schematic of the paper-based microfluidic device design for NoV detection using NoV aptamer functionalized MWCNTs. **c** the picture of the paper-based microfluidic device.

## Experimental

### Materials and chemicals

Carboxylic acid functionalized MWCNTs, graphene oxide, nitrocellulose membranes (Whatman® Protran®) and all other mentioned chemicals and solvents were purchased from Sigma-Aldrich (Oakville, ON, Canada). The 6-carboxyfluorescein (6-FAM) binding norovirus aptamer was synthesized by Integrated DNA Technologies, Inc. (Coralville, IA, USA) with the sequence of 5’-AGT ATA CGT ATT ACC TGC AGC CCA TGT TTT GTA GGT GTA ATA GGT CAT GTT AGG GTT TCT GCG ATA TCT CGG AGA TCT TGC-3’ [20]. The virus like particles (VLPs) formed by group 2 Norovirus capsid antigen were purchased from MyBioSource, Inc. (San Diego, CA, USA). The craft paper punch was purchased from Amazon.ca (McGill®, Paper Blossoms Lever Punch-Multi Daisy, .625" To 1"). Unless otherwise noted, all reagents were of analytical grade unless otherwise stated, all solutions were prepared with double-distilled water.

### Preparation of 6-FAM Aptamer Functionalized and MWCNTs /GO

The probe of 6-FAM aptamer was prepared by firstly resuspending the dried aptamer pellet in the TE buffer (10 mM Tris HCl, 0.1 mM EDTA, pH 8.0) and followed by incubation for 30 min at room temperature. Then a series of aptamer solution was made with folding buffer (1 mM MgCl_2_, 1×PBS, pH 7.4) and heated at 85°C for 5 min. This was then followed by cooling down the dilutions before use.

MWCNTs and GO were diluted with DI water to a series of concentrations ranging from 0.05 to 0.1 mg/mL. Then we mixed the MWCNTs or GO dilutions with 6-FAM aptamer working solutions of specific concentrations and incubated for a period of time to quench the fluorescence of the aptamer. The mixture was then pipetted onto the detection area of the paper-based microfluidic device. The incubation time was optimized by investigating the fluorescence signal at time points of 1 min, 5 min, 10 min and 20 min. Serial dilutions of NoV VLPs stocks (10-fold) were prepared on a range of 0~12.9 μg/mL using 1X phosphate buffered saline (PBS Buffer) solution.

### Paper-based microfluidic device fabrication and assay procedure

The schematic of the paper-based microfluidic device design illustrating the sensing mechanism as well as the picture of the device is shown in Fig. 1c and 1d. A blossoms lever-shaped paper device was patterned manually as the paper-based microfluidic device on the nitrocellulose membranes by a craft punch under room temperature. The width of the “arm” channels was 500 μm, the diameters of the center area (sample loading point) and the detection area were 3 mm and 1 mm, respectively.

Nitrocellulose membranes were used to make the paper-based microfluidic device. Nitrocellulose membranes are a popular matrix that can be used for simple and rapid protein immobilization because of its non-specific affinity for amino acids [21]. The mixture of 6-FAM aptamer solution and MWCNTs or GO was pipetted onto the detection area of the paper-based microfluidic device and allowed to dry in the air for 20 min. The dried paper was then undergone the blocking treatment by using PBS with 5% BSA and 0.05% Tween-20 for 30 min. The blocked paper device was then ready to use and could be stored at 4 °C for short term storage. It is noted that the nitrocellulose membrane based paper device should be kept in a dry atmosphere and away from noxious fumes, avoiding exposure to sunlight, which is conducive to extending the shelf life of nitrocellulose membranes-based diagnostic device and maintaining the molecular recognition capability of immobilized proteins for a long time, even a few years.

During an assay, the fluorescence intensities of the reaction (sensing) zone were recorded as reference. Then aliquots of varying concentrations of NoV VLPs in PBS buffer (10μL) and controls were loaded onto the central, the liquid would diffuse into the detection area spontaneously due to the capillary force. After incubation, the fluorescence intensities of the reaction (sensing) zone were measured again to quantitatively analyse the sample concentration. Each sample was tested 3 times. Three independent experiments were carried out for each conditions.

### Characterization and optimization

Transmission electron micrographs were obtained using the FEI-Tecani G2, operating at 200kV. The fluorescent imaging (Ex/ Em=490nm/520 nm) of the detection area was taken on a fluorescent microscopy (Nikon Eclipse Ti, Nikon Canada Inc., Mississauga, ON, Canada). All images were taken under the same settings, namely exposure time, magnification, etc. The fluorescence intensity was then analyzed by Nikon NIS Elements BR version 4.13 software to investigate the concentration of the sample. For the validation tests, the fluorescence spectra were recorded by the Cytation 5 Multi-mode Reader (BioTek, Winooski, VT, USA).

The ratio of the 6-FAM aptamer solution and MWNCTs/ GO, quenching and recovery time were optimized. In a typical optimization test, 20 μL of 6-FAM aptamer, 20 μL of MWCNTs/ GO solution were well mixed. Fluorescence intensity was measured at the time points of 0, 5 min, 10 min, 15 min and 20 min. Afterwards, 20 μL of NoV VLPs standard solution was added and mixed well. After incubation for a period time, the recovered fluorescence intensity of the resulting solutions was analyzed. The relationship between the fluorescence intensity change and the corresponding concentration was plotted. The fluorescence spectra were measured at Ex/ Em= 490 nm/ 520 nm by the Multi-mode Reader. All samples were prepared in triplicates.

## Results and discussion

### Characterization, validation and optimization of the biosensor

The TEM images of the carboxylic acid functionalized MWCNTs and GO clearly showed the tubular and sheet nanostructures (Fig. 2a and 2b). The UV-Vis absorption spectra of the carboxylic acid functionalized MWCNTs and bare GO are shown in Fig. 2c and as shown the maximum absorption peaks appear at 253 nm and 258 nm, respectively. Fig. 2d shows the fluorescence spectra of the 6-FAM NoV aptamer. The fluorescence intensity peak is located at a wavelength of 520 nm.

**Fig. 2.**
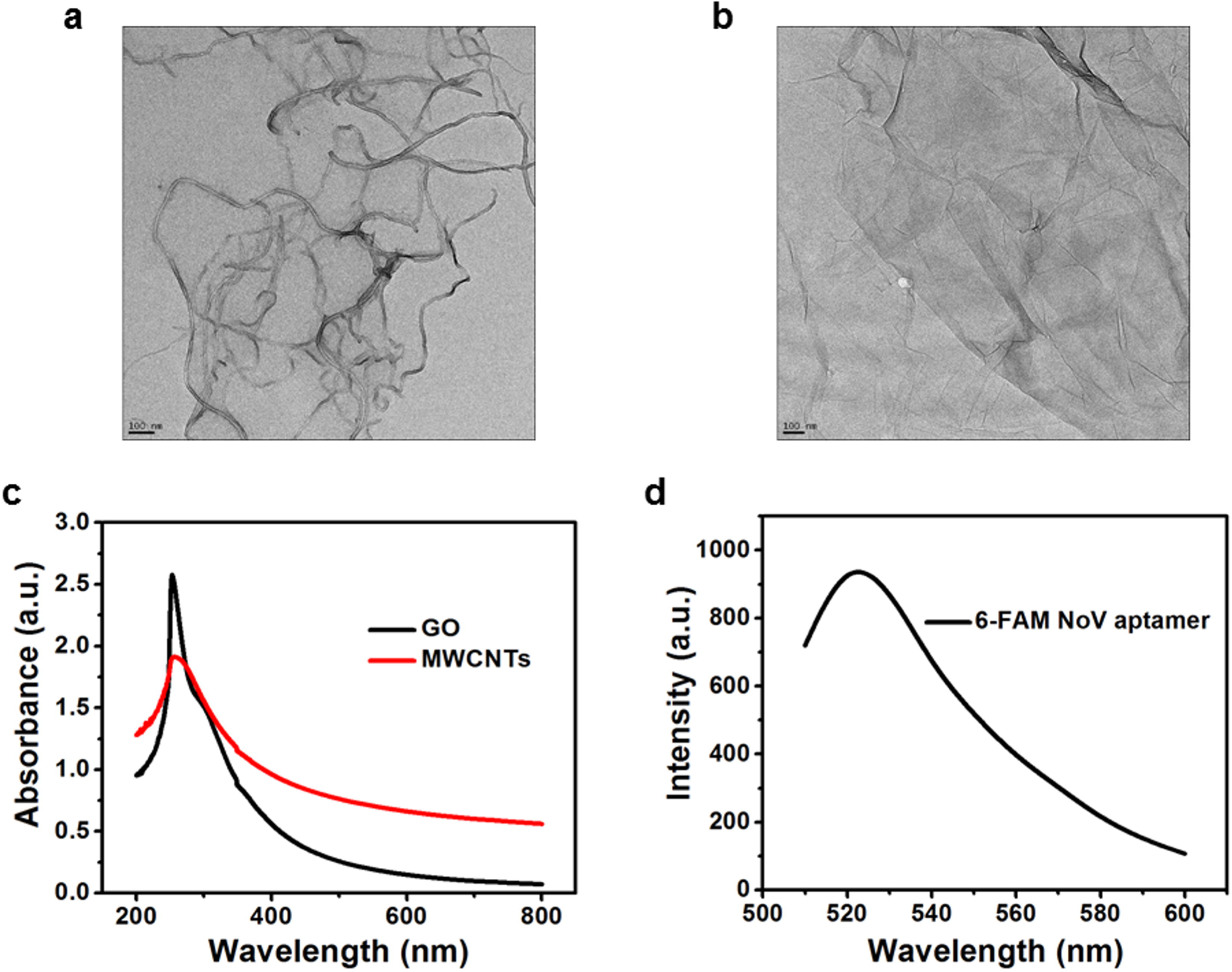
Characterization of the carboxylic acid functionalized MWCNTs and GO. **a** TEM images of MWCNTs. **b** TEM images of GO. **c** UV-Vis absorption spectra of the MWCNTs and bare GO. **d** Fluorescence spectra of 6-FAM NoV aptamer (Ex=490 nm, Em=520 nm)

The quenching effect of two nanomaterials, MWCNTs and GO, were investigated and compared. In order to obtain optimum performance, the concentrations of the MWCNTs, GO, NoV aptamer as well as the reaction (incubation) time were studied, respectively. For the nanomaterials MWCNTs and GO, a series concentrations, 0.05, 0.1, 0.5 and 1.0 mg/mL, were investigated, respectively. And for the NoV aptamer, 1 μM and 5 μM were chosen for further experiments. The influence of reaction time on the fluorescence intensities of before and after quenching were also studied. Fig. 3 and Fig. 4 show the fluorescence quenching and recovery of the NoV aptamer of 1 μM and 5 μM by various of concentrations of MWCNTs and GO when detecting the NoV sample of 129 ng/mL. The validation of these optimization testing was conducted by a microplate reader. All assays were performed at room temperature. As shown in Fig. 3a and 3b, higher quenching effect was found at the 0.1 mg/mL of MWCNTs when NoV aptamer was at 1 μM while 0.05 mg/mL of MWCNTs had higher one when the concentration of NoV aptamer increased to 5 μM. For the nanostructure GO, there is no notable difference between 0.05 mg/mL and 0.1 mg/mL was observed when NoV aptamer was at 1 μM and 5 μM, as shown in Fig. 4a and 4b. A slightly strong quenching effect was found by GO compared to that of MWCNTs, which can be attributed to the more interaction area of the sheet structure of GO than the tube structure of MWCNTs. Fig. 3c, 3d and Fig. 4c, 4d shows the recovering of the fluorescence quenched by MWCNTs and GO, respectively. Remarkable recovery of the fluorescence quenched appeared for both of them, which was attributed to FRET generated by the synergy of particles collision and π-π stacking interaction between them and aptamer [22]. However, the recovery level varies. For the MWCNTs, the optimum recovery appeared at the setting of NoV aptamer of 5 μM and MWCNTs of 0.1 mg/mL, as shown in Fig.3d. While for the GO, the optimum recovery appeared at the setting of NoV aptamer of 5 μM and GO of 0.05 mg/mL. A slightly strong quenching effect was found by GO compared to that of MWCNTs, which can be attributed to the more interaction area of the sheet structure of GO than the tube structure of MWCNTs. Thus a relative recovery rate can be achieved by GO.

**Fig. 3.**
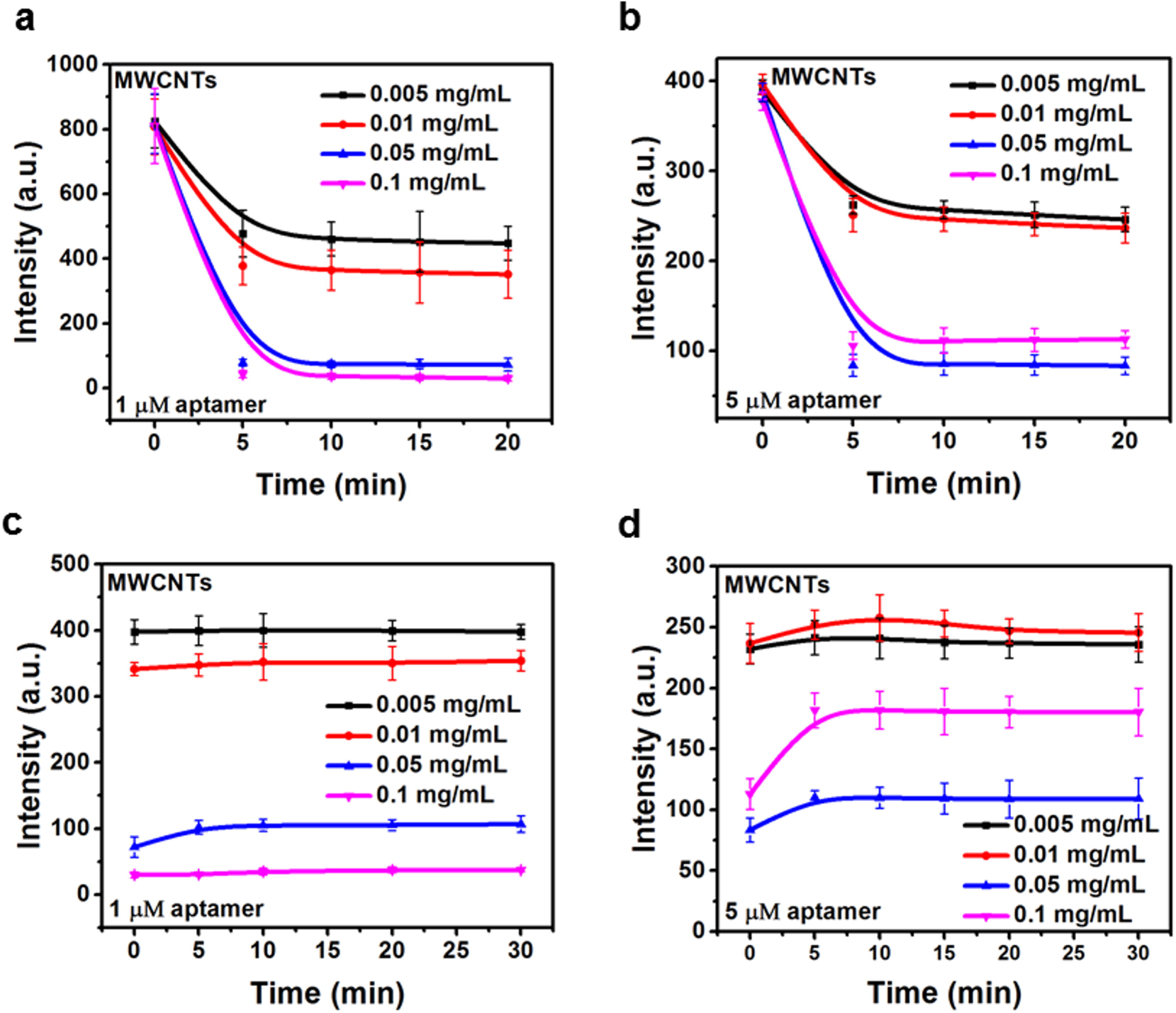
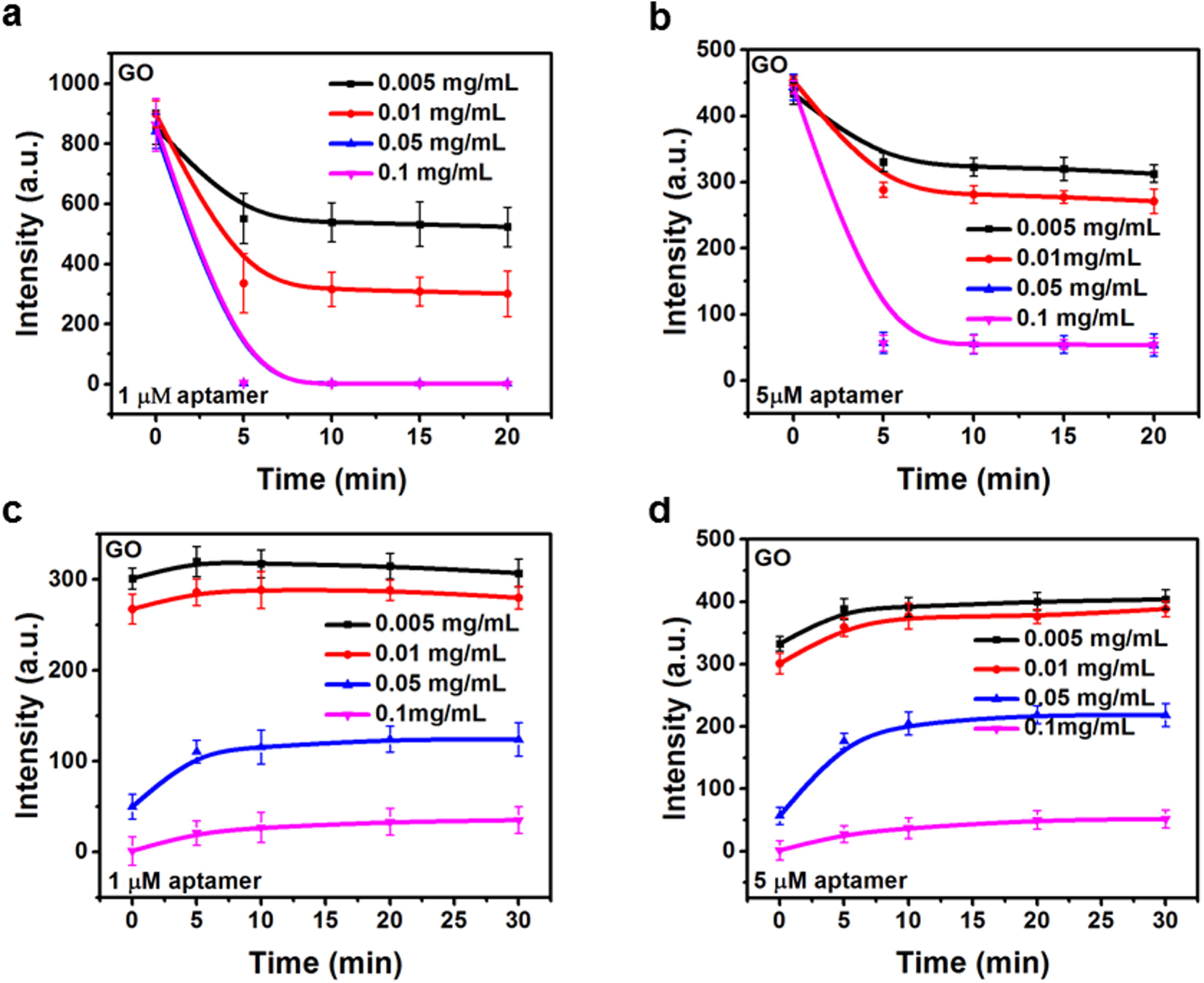
Quenching effect of MWCNTs of various concentrations at **a** 1 μM and **b** 5 μM NoV aptamer versus time. Recovery effect of MWCNTs of various concentrations at **c** 1 μM and **d** 5 μM NoV aptamer versus time.

**Fig. 4.**
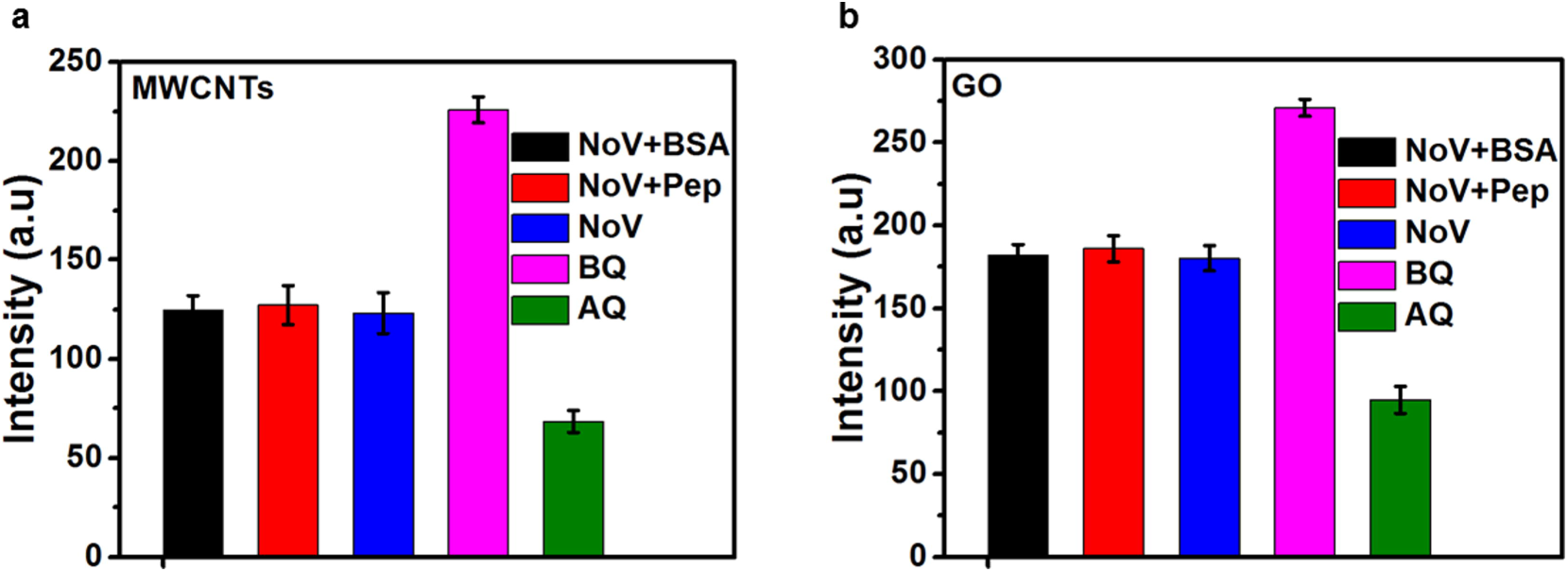
Quenching effect of GO of various concentrations at **a** 1 μM and **b** 5 μM NoV aptamer versus time. Recovery effect of GO of various concentrations at **c** 1 μM and **d** 5 μM NoV aptamer versus time.

Fig. 3 and Fig. 4 also show the time-dependent fluorescence intensity changes. The time-dependent response was studied to obtain a suitable incubation and reaction time for quenching and recovery. As shown in the figures, the quenching of the 6-FAM NoV aptamer occurred immediately after adding MWCNTs or GO, and a dramatic decline was observed within 5 min, and then became a plateau after prolonging time. As indicated in the Fig. 3 and Fig. 4, the recovery of the fluorescence could be completed within 5 min, because no notable intensity changes appeared afterwards. Thus 5 min was considered as the optimal incubation time for quenching as well as the optimal recovery time.

### Specificity of the biosensor

We evaluated the selectivity of our biosensor by detected the NoV VLPs solution mixed with bovine serum albumin (BSA) and peptidoglycan, respectively. BSA was used as a blocking buffer for the paper based device, while peptidoglycan is the major cell wall component of both the gram positive and gram negative bacteria and may present in complex food matrices. The mixture of BSA and NoV VLPs (3.3 mg/mL and 129 ng/mL in final concentration), and the mixture of peptidoglycan and NoV VLPs (1.2 mg/mL and 129 ng/mL in final concentration) were detected. The results as shown in Fig. 5 confirms the excellent specificity of the developed biosensor. Notable changes in recovered fluorescence intensity were observed in all solutions due to the presence of NoV VLPs. However, no distinguishable changes in recovered fluorescence intensity were observed among the BSA or the peptidoglycan mixture with the NoV VLPs solution indicating the absence of any cross-reactivity. The results also demonstrate that the selected NoV aptamer can only specifically bind to the NoV VLPs.

**Fig. 5.**
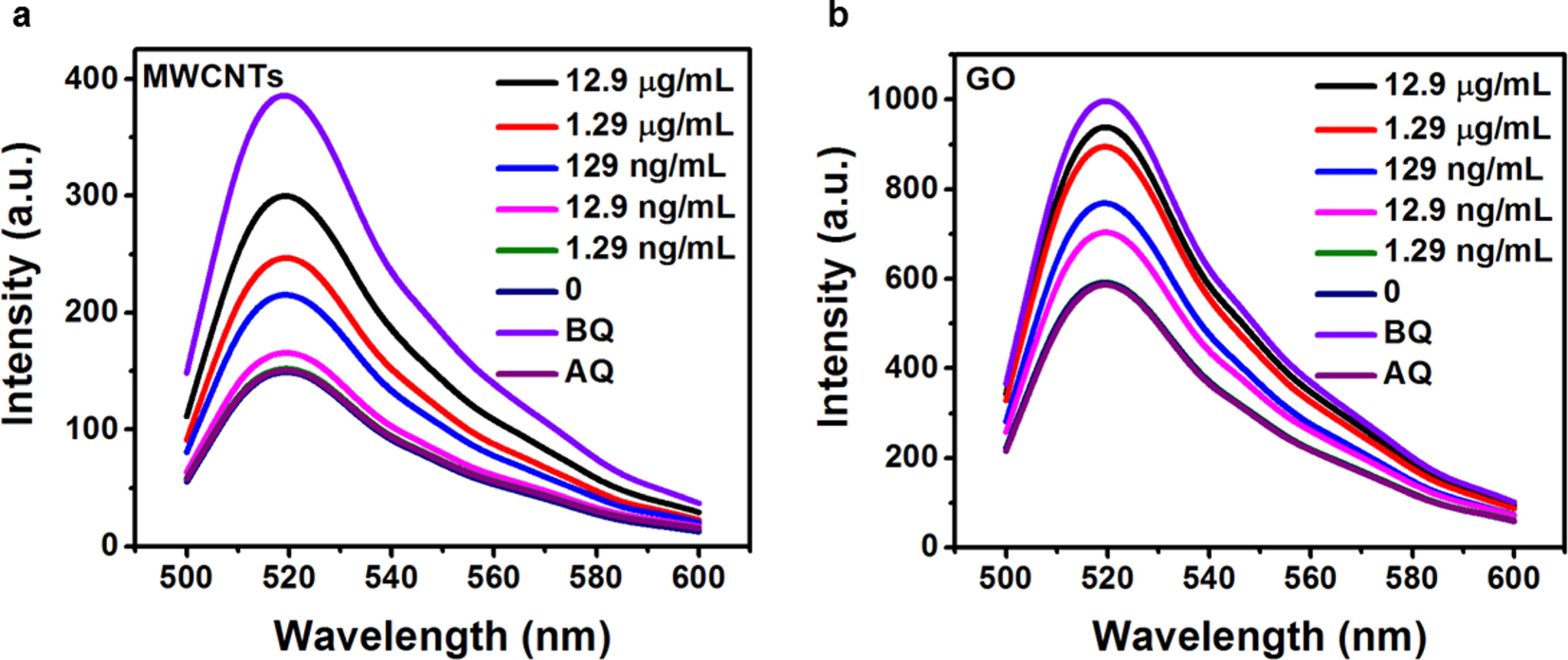
Investigation of the selectivity of the biosensor by testing NoV VLPs sample with BSA and peptidoglycan. **a** MWCNTs **b** GO.

### Detection of NoV VLPs

As a validation, a series of NoV VLPs dilution were detected by a microplate reader under the optimized parameters. Fig. 6 shows the recovered fluorescence spectra versus various concentration of NoV VLPs ranging from 0 to 12.9 μg/mL. The recovered fluorescence intensity increase with respect to the elevated protein concentration. No significant difference was observed between the spectra of 1.29 ng/mL and that of after quenching. However, distinguishable fluorescence intensity changes appear when the concentration increased to 12.9 ng/mL for both the nanomaterials.

**Fig. 6.**
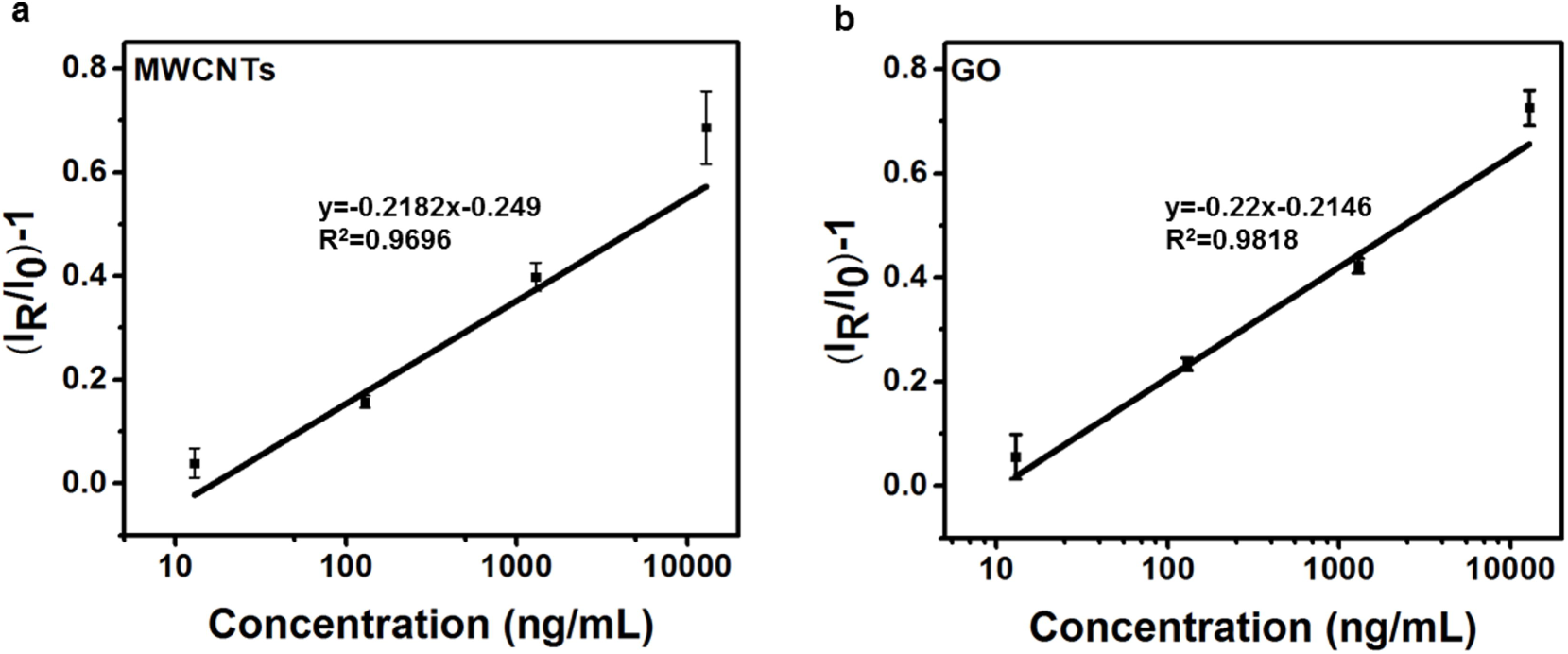
Fluorescence spectra of the 10-fold NoV VLPs serial dilution of NoV VLPs in buffer ranging from 0 to 12.9 μg/mL by **a** MWCNTs **b** GO.

Under the optimized conditions, a series of NoV VLPs dilutions were assayed on the paper-based microfluidic device. For multiplex detection, up to 5 sample dilutions and a negative control could be assayed on a ready-to-use paper-based microfluidic device at one time. All tests were performed at room temperature. To reduce the errors, the difference between the recovered fluorescence intensity (I_R_) and after quenching (I_0_) was used to determine the NoV VLPs concentration in the sample. Fig. 7 shows the calibration curve of NoV VLPs assay by plotting the changes in fluorescence intensity versus the different concentration, with the linear range of 0~12.9 μg/mL and R^2^ being 0.9696 for MWCNTs and 0.9818 for GO. The LoD is calculated by kS_b_ S [23], where S_b_ is the standard deviation of the blank measure, S is the sensitivity (Δconcentration/Δintensity) and k = 3 is numerical factor. The LoDs of the paper-based microfluidic biosensor are 40 pM for MWCNTs and 30 pM for GO with 3% −7% in relative standard deviation (RSD).

**Fig. 7.**
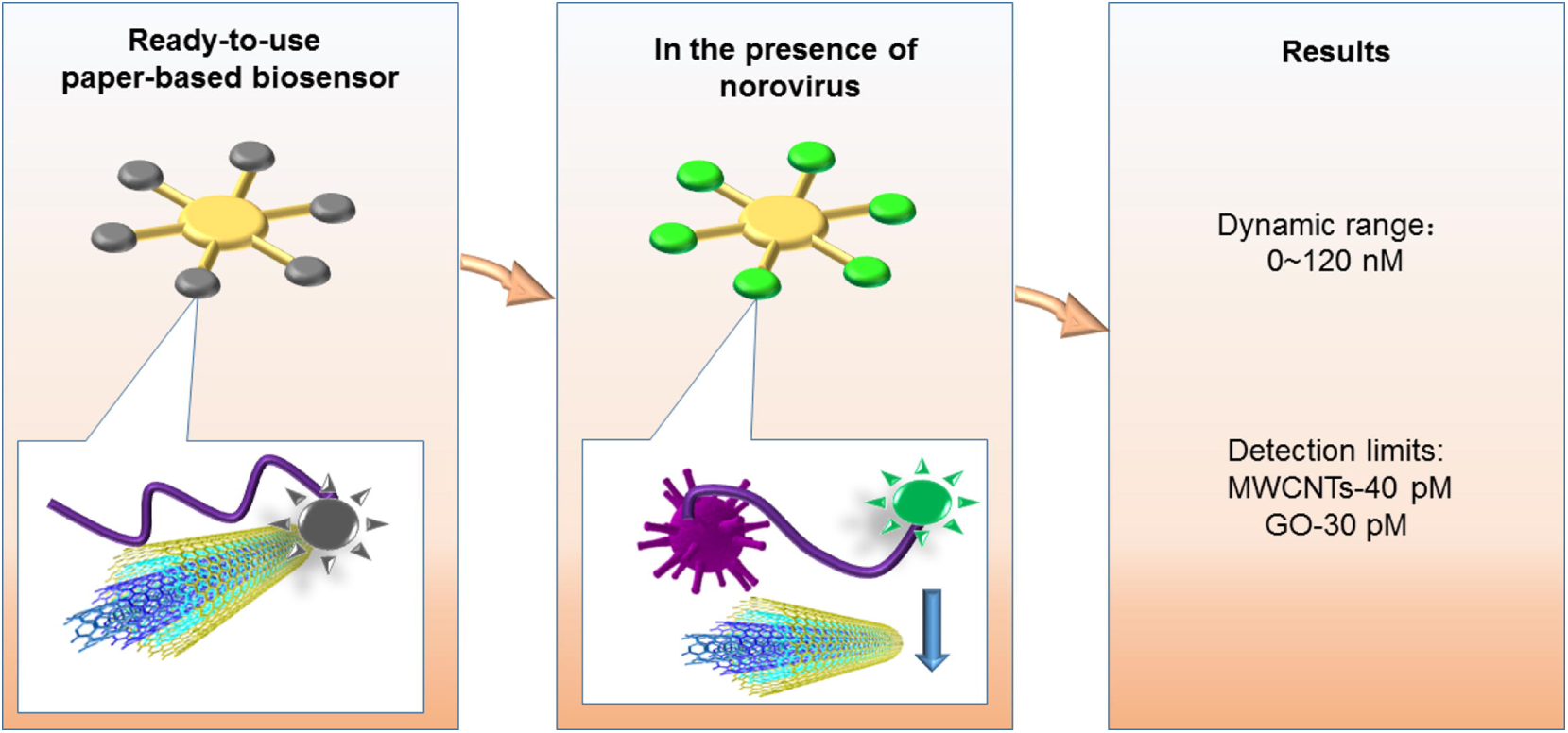
Standard curves of the NoV VLPs detection by **a** MWCNTs. **b** GO on the paper-based microfluidic device via the fluorescence intensity changes versus various NoV VLPs concentration ranging from 0 ~ 12.9 μg/mL.

## Conclusions

A paper-based microfluidic biosensor is developed for the detection of norovirus. The biosensor utilized the FRET between the 6-carboxyfluorescein labelled norovirus aptamer and MWCNTs and GO. The MWCNTs would quench the fluorescence of aptamer when they were mixed together. In the presence of NoV, the fluorescence would be turned on (recovered) due to the specific binding with the aptamer. The quantitative results showed a limit of detection of 40 pM for MWCNTs and 30 pM for GO, respectively. The easy-to-make paper-based microfluidic biosensor could be easily applied for multi-virus diagnosis hence finds numerous applications in food sector and clinical diagnosis.

## Abbreviations

NoV: noroviruses
6-FAM: 6-carboxyfluorescein
MWCNTs: multi-walled carbon nanotubes
GO: graphene oxide
FRET: fluorescence resonance energy transfer
NoV VLPs: norovirus virus-like particles
ELISA: enzyme-linked immunosorbent assay
RT-PCR: reverse transcription polymerase chain reaction
LoD: limit of detection
LFAs: lateral-flow assays
POCT: point-of-care testing

## Authors’ contributions

XW and SN designed the study; XW performed experiments, acquired and analyzed data, XW and SN drafted and edited the manuscript. All authors read and approved the final manuscript.

## Acknowledgements

The authors sincerely thank the Natural Sciences and Engineering Research Council of Canada (400705) for funding this study.

